# Characterization and analysis of neuronal signaling using microelectrode array combined with rapid and localized cooling device for cryo-neuromodulation

**DOI:** 10.1101/2024.08.16.608362

**Authors:** Jaehyun Kim, Jong Seung Lee, Soyeon Noh, Eunseok Seo, Jungchul Lee, Taesung Kim, Seung-Woo Cho, Gunho Kim, Sung Soo Kim, Jungyul Park

## Abstract

Cryoanesthesia––a purely physical anesthesia treatment that freezes tissue and attenuate nerve activity––can provide fast treatment through freezing and thawing of cryo-machine and is inexpensive compared to other anesthetics. However, cryoanesthesia has not been widely adopted because securing safe and effective conditions requires quantitative measurement and analysis of neuronal signaling during freezing and recovery, for which research tools are limited. A lack of rapid and localized cooling technologies for quantitative cellular level analysis, in particular, hinders research on not only the optimal cryo-modulation of neuronal activities but also its influence to neighboring cells via cellular networks. Here, we introduce a novel cryo-neuromodulation platform, a high-speed precision probe-type cooling device (∼20°C/s at cooling) that provides localized cooling combined with a microelectrode array (MEA) system. We explored the temperature conditions for efficient silencing and recovery of neuronal activities without cell damage. We found that electrical activities of neurons were fully recovered within 1 minute of cooling duration with the maximum cooling speed, which was also confirmed with calcium imaging. The impact of silenced neurons on the neighboring neural network was explored using the localized cooling and we perceived that its influence can be transmitted if the neuronal network is well organized. Our new cryo-device provides rapid and reversible control of neural activities, which allows not just quantitative analysis of the network dynamics, but also new applications in clinical settings.

## Introduction

Cryoanesthesia, which does not use chemicals but a pure physical principle, is efficient in reducing pain, provides fast treatment^1^, is less expensive than other topical anesthetics^2^, and thus has been used in cataract surgery, acne treatments, dental treatment^1, 3, 4^, treating or killing cancer cells^5, 6, 7^, etc. However, it has been doubtful to guarantee safe anesthesia. In lab environments, several approaches have been used to study the cold tolerance and behaviors of cells and tissues at low temperatures, such as baths^8^, chambers^9, 10^, or peltiers^11, 12, 13, 14, 15, 16^. However, they are difficult to control the temperature precisely and their dynamics is not fast, preventing neurons from safe recovery^17^. Furthermore, the size of current cooling devices is too large for precise spatial modulation, limiting their applications to study single cell level or signal transmission between neurons. Other recent neuromodulation techniques, such as optical neuromodulation^18, 19^, thermal control neuromodualtion^20, 21, 22^ by using nanomaterials, and magnetic neuromodulation^23, 24^, can control neuronal signaling at the cellular and tissue level. Particularly, Yoo *et al*.^21, 22^ have achieved reversible and localized neuromodulation without cell damage using gold nanorod, which enables rapid increase in local temperature with photothermal devices on microelectrode array (MEA). However, due to the lack of cooling methods that enable precise and fast decrease in local temperature, there is, as far as we are aware, no demonstration to show safe and reversible cellular level cryo-neuromodulation.

This study proposes a cryo-neuromodulation technology that combines a rapid and accurate cooling device for localized cooling with ∼20°C/s, which addresses critical limitations of existing cooling technologies, and MEA, which measures microcurrents at the single-cell scale (Fig. 1). We explored the parameter space of the cooling temperature, duration, and speed to discover conditions for safely manipulating neuronal signaling. We measured the health of neurons by comparing inter-spike interval, spike rate, spike amplitude, inter-burst interval, and spike width before and after cooling^25, 26, 27, 28, 29, 30^. The effect of localized cooling on neighboring neural networks was also studied, and it was transferred to long-range cells when the networks were well organized. This new cryo-neuromodulation device can serve as a fundamental experimental platform to provide rapid, safe, reversible anesthesia. Furthermore, it has a potential to be used in therapeutic approaches where neuronal signaling control at low temperatures is beneficial.

**Fig. 1.**
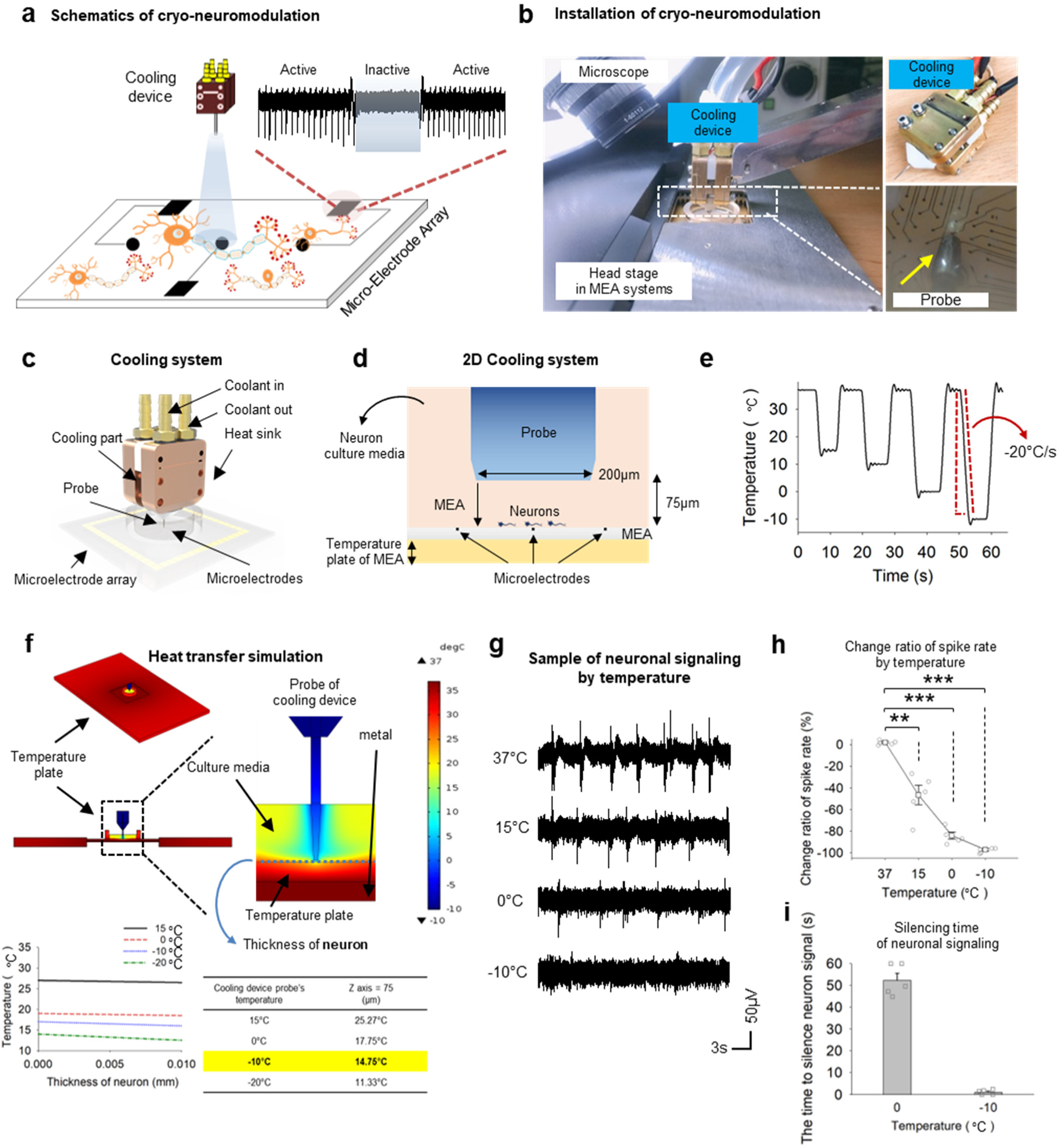
In vitro cryo-neuromodulation system enabling spatially localized rapid cooling. (a) Schematics of cryo-neuromodulation system: The neurons cultured on the MEA are cooled by the rapid and localized cooling device. (b) Equipment configuration: A probe-type cooling device is positioned close to neurons on top of MEA device, under a CCD-camera equipped microscope. (c) The cooling system consists of a probe with a copper bonded to thermoelectric cooler (TEC) in the cooling part and heat sinks to dissipate heat. (see Methods for details) (d) 2D MEA cooling system. Thermoelectrically cooled probe reduces the temperature of neurons on MEA chip and potential changes of the neuron detect by microelectrodes on the MEA chip. (e) The performance of cooling device. It has maximum velocity -20℃/s. (f) Top left: dark red square is temperature plate, and dotted square are the probe of the cooling device and MEA filled with culture media. Top right: blue dot line is thickness of neurons. Down left: following thickness of neuron, the actual temperature distribution. Down right: Temperature of cooling device and actual temperature at z coordinate 75 µm. (See Extended Data Fig. 2 in detail) (f) Neuron signals measured by the MEA at each temperature. (g) The firing rate and pattern of action potentials significantly changes according to temperatures. (paired t-test: 37℃ vs 15 p = 0.0012 with n=5, 37℃ vs 0℃ p < 0.001 with n=5, 37℃ vs -10℃ p value < 0.001 with n=5). (h) Change ratio of spike rate by temperature. (Change ratio of spike rate: [(Spike rate (after cooling) – Spike rate (before cooling)) / spike rate (after cooling)] * 100) (i) The time to silence neuron signal at each temperature. (one sample t-test: mean time 1.2 vs times at -10℃ p = 0.9402 with n = 5, mean time 52.3 vs times at 0℃ p = 0.9786 with n = 5). * p<0.05, **p<0.01, ***p<0.001

## Results

### The new cryo-neuromodulation system enables fast and spatially focused temperature modulation

In most cooling experiments, the entire bathing media is cooled, slowing down the process (below 2℃/s^25, 31, 32, 33, 34, 35^). On the other hand, our device is equipped with a fine probe to enable spatially focused temperature modulation (Fig. 1), and the temperature of the probe can be modulated fast: Reaching from 37℃ to -10℃ takes an average of 2.8 seconds (-20℃/s; Fig. 1e), allowing fast localized cooling. With this device, we first identified the probe temperature at which the neurons in the medium under the probe were silenced. We located the cooling device 50 ∼100µm above the cultured neurons on the MEA and measured the normal neural activity for 3 to 5 minutes at 37℃. We then lowered the temperature of the cooling device to 15℃, 0℃, and -10℃ (Fig. 1a and g). At 37℃, spikes were clearly observed, but the spike rate was gradually decreased at 15℃ (50%), 0℃ (85%) and eventually no spike was observed with the probe at -10℃ (100%; Fig. 1h)^33, 36^. The time to silence neuron signal depends on the cooling temperature; 55 seconds at 0℃ and 5 seconds at -10℃ (Fig. 1i). Note that, in multiphysics simulation, the temperature around the neurons was estimated to be 15.82°C with -10°C probe temperature (Fig. 1f)^37^, which was consistent with other studies showing silenced neural activities at 15∼20℃^37, 38^.

### Neuronal activity recovers better with a shorter cooling duration

A long exposure to the low temperature may impose stress on neural tissues, resulting in an incomplete recovery or even damage to the circuits. Our device allows precise control of the cooling duration, allowing us to evaluate the effect of the cooling duration on the recovery of the neural activity (Fig. 2). The cooling duration was set to 1, 3, 5, and 10 minutes (Fig. 2 and Supplementary. Fig. 5), and the neuronal signals before and after cooling to 15℃ (probe temperature at -10℃) were compared. We found that the time for the first spike reappearance after the end of cooling was not significantly different between 1 and 5-minute (< 5 s) but increased with 10-minute cooling duration (13 s) (Fig. 2d). The signal recovery time (see Methods) was much longer than that of the first spike, which reached 40 seconds for the 10-minute cooling duration. The difference of spike amplitude before and after cooling was not significant for 1-minute cooling but was prominent for 10-minute cooling (Fig. 2e). We also observed an insignificant change in the firing rate for the 1-minute cooling duration, but a significant change for the 10-minute duration (decreased inter-burst-interval (IBI) times than before cooling) (Fig. 2f). Specifically, IBI was considerably reduced by longer cooling duration^39, 40^ (Fig. 2g-h), potentially reflecting the reduced number of neurons participating in bursting activity. On the other hand, the inter-spike-interval (ISI) was increased for the 10-minute cooling duration (Fig. 2i-k), indicating slower neural dynamics. Finally, the spike width was significantly increased for 10-minute cooling duration (Fig. 2l-m) in line with a recent report^11^. Overall, we concluded that neurons could not fully recover after the 10 minutes of cooling, suggesting significant damage or deteriorating effect to the neural dynamics.

**Fig. 2.**
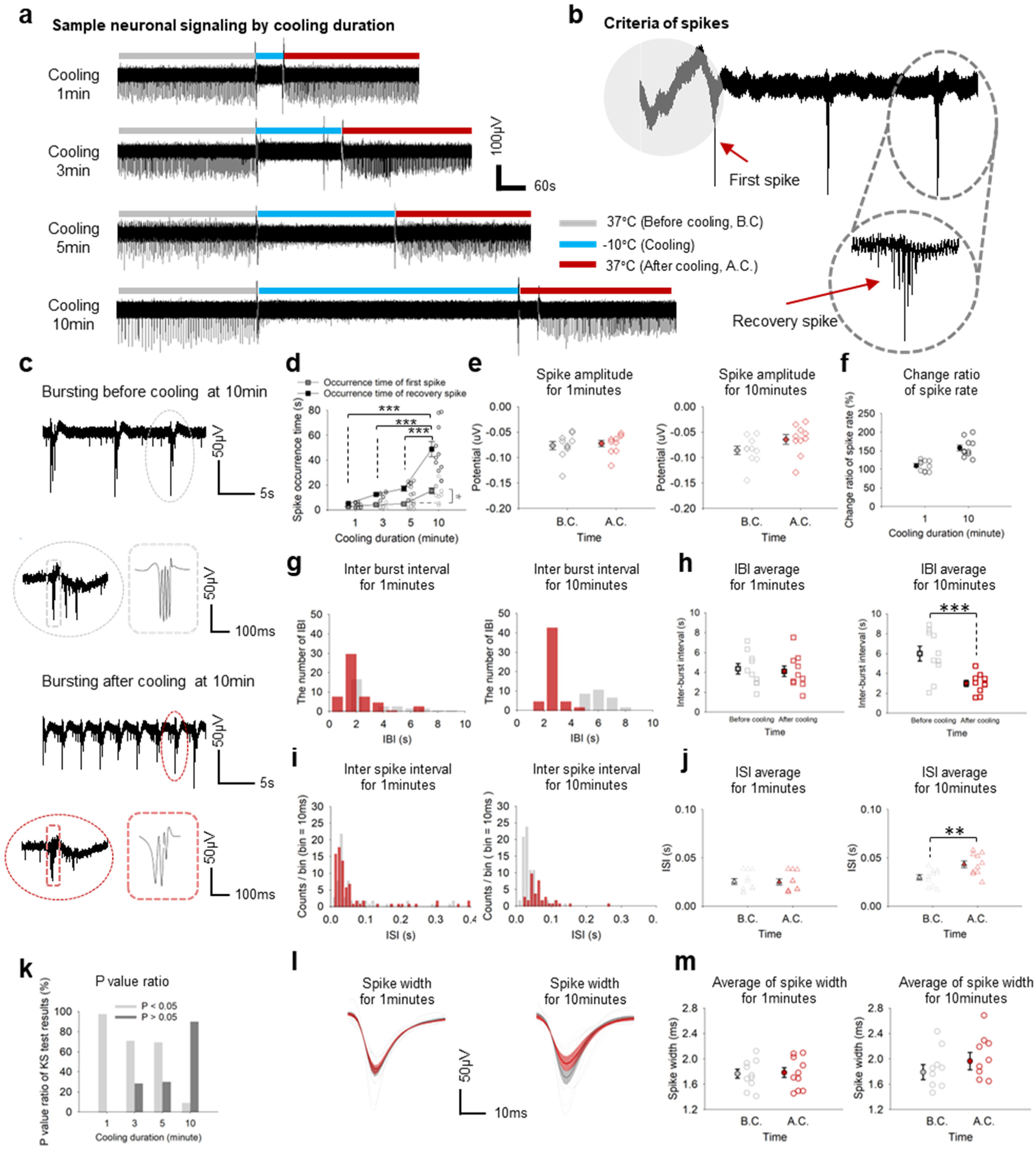
The silencing and recovery of neural activity depends on the duration of cooling. (a) Activity of a sample neuron with respect to different cooling durations. (b) The criteria of first spike and recovery spike. (c) Bursting before cooling at 10minutes; Top: neural signal sample for 15 seconds. Down left: Bursting in gray circle dot line of top figure enlarged. Down right: Part of bursting in gray square dot line expanded as simple figure. Bursting after cooling at 10minutes; Top: neural signal sample on MEA for 15 seconds. Down left: Bursting in red circle dot line of top figure enlarged. Down right: Part of bursting in red square dot line expanded as simple figure. (d) Occurrence time of the first spike and the recovery spike according to different cooling durations. (For the first spike paired t-test; 1 minute vs 3 minutes, n=10, p=0.1989: vs 5 minutes, n=10, p<0.05: vs 10 minutes, n = 10, p<0.05), (For the recovery spike paired t-test; 1 minute vs 3 minutes, n=10, p<0.001: vs 5minutes, n = 10, p<0.001: vs 10 minutes, n=10, p<0.001). (e) Comparison of spike amplitude before and after cooling Left: 1-minute cooling duration (paired t-test: Before cooling vs After cooling, n=10, p=0.6854) right: 10-minutes cooling duration (paired t-test: Before cooling vs After cooling, n=10, p=0.1092) (f) Comparing change ratio of spike rate at 1 minute and 10 minute. (g) Left: The number of IBI histogram at 1-minutes Right: the number of IBI histogram at 10-minute. (h) Scatter plot of inter-burst interval before and after cooling. Left: At 1-minute (paired t-test: Before cooling vs After cooling, n=10, p=0.7311). Right: At 10-minute (paired t-test: Before cooling vs After cooling, n=5, p<0.001). (i) Inter-spike interval histogram; Left: At 1-minute. Right: At 10-minute. (j) Scatter plot of ISI’s average Left: At 1-minute (paired t-test: Before cooling vs after cooling, n=10, p=0.9016). Right: At 10-minute (paired t-test: Before cooling vs after cooling, n=10, p<0.01) (k) P-value of KS-test result by cooling duration. If P value is larger than 0.05, before and after cooling have different distribution. On the other hand, If P value is lower than 0.05 before and after cooling have identical distribution. (l) Average and Standard error spike width. (see Methods) Left: At 1-minute. Right: At 10-minute. (m) Scatter plot for average of spike width before and after cooling Left: At 1-minute (paired t-test: Before cooling vs after cooling, n=10, p=0.857) Right: At 10-minutes (paired t-test: Before cooling vs after cooling, n=10, p value=0.3487). (See Extended Data Fig. 3 in detail at 3, 5-minutes). * p<0.05, **p<0.01, ***p<0.001

### Faster cooling speed allows better recovery

Conservation of biological tissue benefits from fast cooling^41, 42^. However, the cooling speed of most commercial devices (or perfusion with cooled solution) has been slow^25, 31, 35, 43, 44, 45, 46^. Our probe provides unprecedented control over the cooling speed with high spatial specificity. We compared the recovery of neural activity after two different cooling speeds: -20℃/s and -4℃/s (Fig. 3a-b). The time to reach the target temperature at maximum speed (-20℃/s) was 2.73 seconds on average, a considerably shorter than 10.33 seconds at -4℃/s. We repeated the sequence of 1 min of cold temperature and 2 minutes of 37℃. The spike amplitude significantly decreased when the speed was slow at -4℃/s (paired t-test between baseline, *i.e.,* before cooling, and trial 5, p=0.0236) whereas it did not significantly change up to 5th repetition with the speed of -20℃/s (Fig. 3c). The IBI decreased with -20℃/s only after the 6th repetition, whereas, at -4℃/s, there was a significant decrease from 4th trial (Fig. 3d) (paired t-test between baseline and trial 4, p=0.0098, between baseline and trial 5, p=0.00062). The spike width did not significantly vary across two cooling speeds (Fig. 3e-f). The ISI significantly increased after repeated cooling but only from 5th trial with -20°C/s whereas from 3rd trial with -4°C/s (Fig. 3g). The ISI was identical up to on average 4 times of trial with -20°C/s, whereas up to on average 2.8 times with -4°C/s (Fig. 3h; Kolmogorov-Smirnov test). These results strongly suggest that neurons are significantly less damaged with, and thus benefit from, faster cooling speed.

**Fig. 3.**
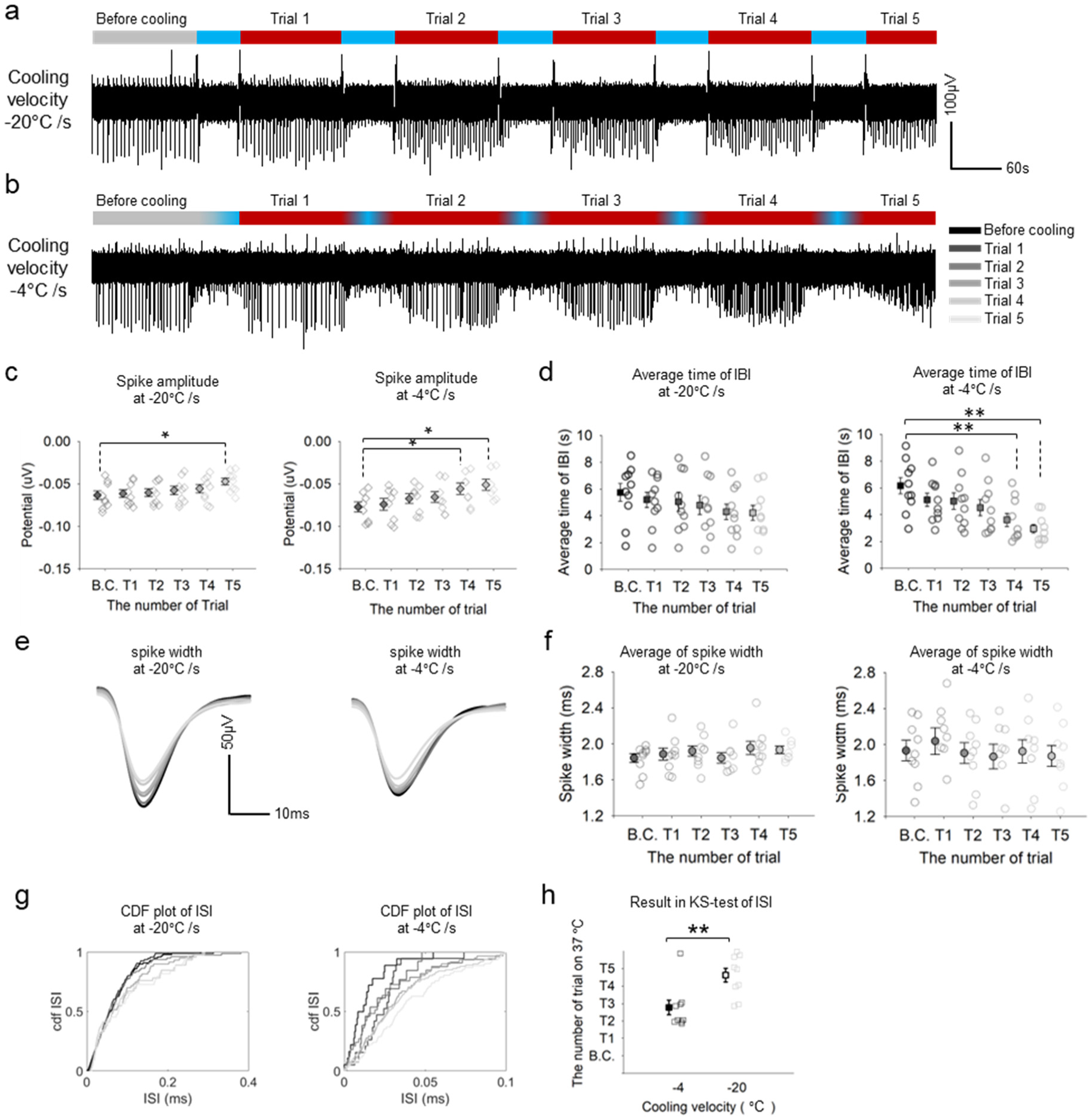
Analysis of neural activity affected by cooling velocity. (a) When cooling velocity is -20°C/s, spike train which was repeated decreasing temperature and increasing. (b) When cooling velocity is -4°C/s, spike train which was repeated decreasing temperature and increasing. (c) Amplitude of neurons spikes by repeating cooling and restoring temperature. Left: At cooling velocity -20°C/s. Right: At cooling velocity -4°C/s. (paired t-test; see supplement table) (d) Average time of IBI by repeating cooling and restoring temperature. Left: At cooling velocity -20°C/s. Right: At cooling velocity -4°C/s. (paired t-test; see supplement table) (e) Spike shape by repeating cooling and restoring temperature. Left: At cooling velocity -20°C/s. Right: At cooling velocity -4°C/s. (f) Spike width by repeating cooling and restoring temperature. Left: At cooling velocity -20°C/s. Right: At cooling velocity -4°C/s. (paired t-test; see supplement table) (g) Cumulative distribution function of ISI Left: At cooling velocity -20°C/s. Right: At cooling velocity -4°C/s. (h) Result in KS-test of comparing ISI distribution before and after cooling. In other words, the number of repetitions until ISI distribution was not equal at cooling velocity -20°C/s and -4°C/s (paired t test; cooling velocity -4°C/s vs -20°C/s, n=10, p<0.01). (See Extended Data Fig. 6 in detail for extra t-test result) * p<0.05, **p<0.01, ***p<0.001.

### Locally silencing neurons reduces network activities far from the cooling site

Silencing neurons in a small area may have impact on the activity of a neural network. To investigate this, we combined our cryo-neuromodulation system with calcium imaging (Fig. 4) ^31, 47^. We first recorded the calcium signal (using Fluo-4) of three neurons near the probe (27µm, 51µm and 103µm away from the probe) before and after cooling. When the cooling began (between 2 and 3 s), the calcium signal rapidly dropped (Fig. 4b and c; more than 80% drop, Fig. 4f). Conversely, the calcium influx was recovered when temperature was restored to a normal temperature. While the neuronal signals were recovered close to the initial status in MEA experiment, the calcium intensity after recovery was much lower than before cooling. This difference may come from the photo bleaching. By setting the bleaching guideline (Fig. 4b) and drawing new plots (Fig. 4d and e) in the controlled experiment, the intensity value was recovered to the before-cooling level or even higher (potentially due to rhythmic rebound activity after inhibition^21, 48^), suggesting full recovery.

**Fig. 4.**
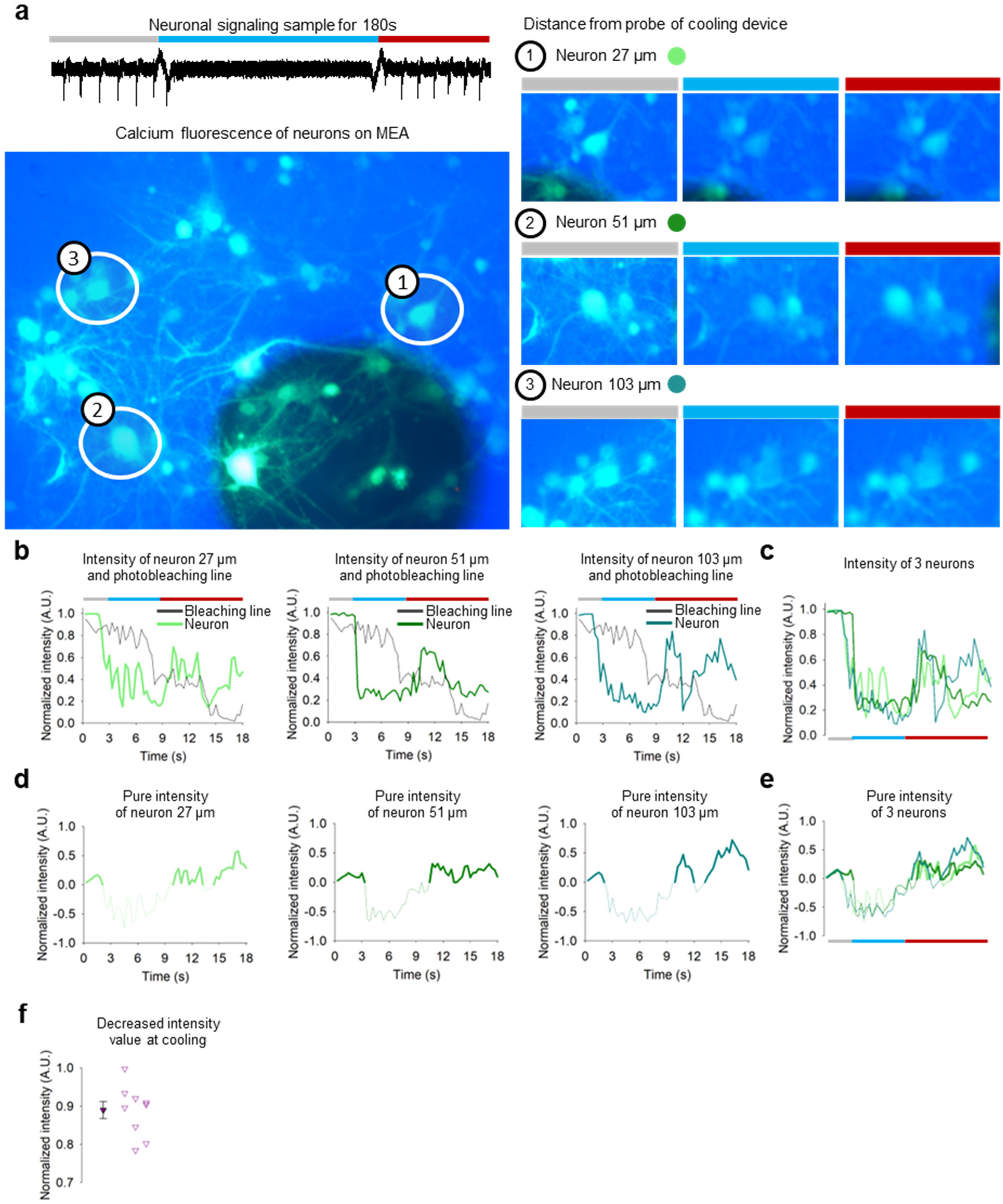
Visualization of the cooling effect to neural activity using calcium imaging. (a) Left top: MEA signal from a neuron before (gray), during (blue), and after (red) cooling. Left bottom: A snapshot of calcium imaging. The shadow of the probe is shown as a big dark circle at the bottom right corner of the image. Right: Calcium imaging snapshots of neurons at 27, 51, 103 µm distance from the cooling point. (b) Normalized intensity of each neurons’ fluorescence with photobleaching guideline. Left: 27 µm Center: 51 µm Left: 103 µm (d) The plot of the intensity of neuron subtracted the photobleaching guideline. The bold line is a positive intensity, and the thin line is a negative intensity. Left: 27 µm Center: 51 µm Right: 103 µm (e) Comparison plot of intensity of neuron subtracted photobleaching guideline for 3 cases. (f) Maximum subtracted intensity before cooling and minimum intensity at cooling. (one sample t-test; the decreased values of intensity vs normalized intensity 0.89, n=10, p value=0.09985)

Next, **t**o investigate the effect of cooling with the distance from the cooling site, neurons were cultured on the whole area of the MEA^21, 49^ (Supplementary Fig. 3). We focused on our experiments for 1-minute cooling duration since longer durations does not guarantee full recovery of neural activity (Fig. 2). The electrode positions were divided into four concentric areas around the cooling probe (Fig. 5a). Naturally, the cooling effect was stronger in a region close to the probe (Fig. 5b): The average firing rate dropped 90% for neurons in the area closest to the probe, whereas neurons showed significantly less effect cooling as the distance increased (-85%, -75% and -40% respectively with increasing distance; paired t-test; Fig. 5b). Furthermore, the average time to silence neurons were higher with the distance (e.g., in the farthest area, most neurons were still active until the end of 1 minute period, Fig. 5c-e). In our simulation and experiments (Fig. 1f, Supplementary Fig. 6-8), the temperature of the closest area from the probe was 15℃ and rapidly increased up to 36℃ with the distance from the probe. This means that, we can silence neurons only in the closest areas (less than 400 µm) by the direct cooling, but not beyond them. However, we observed silencing effect in 400-600 µm distances and occasionally in 600-800 µm distances as well, implying that the silencing neurons in the closest area may have indirect impact on the activity of neurons farther away from the probe presumably via network dynamics. To further investigate this possibility, we carried out a correlation analysis among the active neurons in MEA, to infer functional connectivity in neuronal networks (Supplementary Fig. 10-11); We consider higher correlated neuron signaling from the two different active channels show stronger connectivity between them. When the neuronal signaling in all active channels were silenced by cooling stimulus, the correlation between cooling point and other active channels was high, with an average of 0.8 (Supplementary Fig. 10a). In addition, the average correlation between different channels was also high at 0.8 (Supplementary Fig. 10b). However, when neuronal signaling was silenced only in some portion of channels, the average correlation between the cooling point and other channels was relatively low at 0.68, and some channel showed a correlation of 0.4 (Supplementary Fig. 11a). Moreover, the average correlation between different channels was 0.68. Finally, 16% of channels before cooling and 36% of after cooling showed a correlation of 0.5 or less (Supplementary Fig. 11b). These results suggest that cooling effect can be cascaded onto neighboring neurons via functional connectivity.

**Fig. 5.**
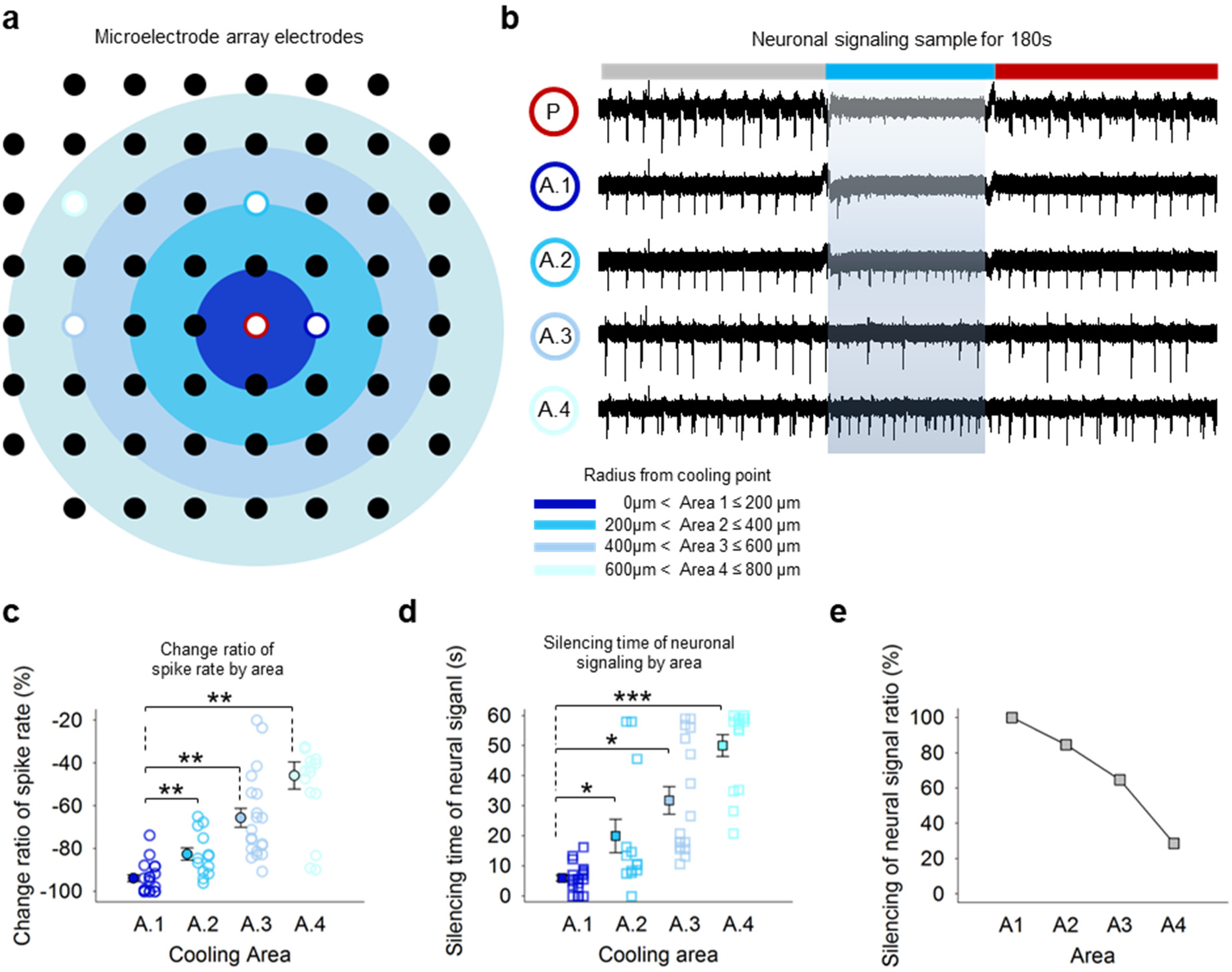
The effect of distance from the cooling probe on neural activity. (a) From the cooling point (red line circle), it is divided into four areas (Area 1 to 4) in a circular shape every 200 micrometers. Each black dot denotes electrodes on MEA, while white dot of blue colors circles each representative electrodes of area. (b) spike train at each electrode. P; the electrode at cooling point, Area1-4; the electrode at area1– 4. (c) Spike rate change across distances (paired t-test compared to Area1; Area2, n=10, p-value < 0.01; Area 3, n=10, p-value < 0.01; Area4, n-10, p-value < 0.01) (d) Silencing time (paired t-test compared to Area 1; Area 2, n=10, p-value=0.0416; Area 3, n=10, p-value < 0.001) (e) Silencing ratio of neuronal signaling by Area.

## Discussion

Our cooling device can rapidly and reversibly inhibit cultured mammalian neurons *in situ*, which we call cryo-neuromodulation. Since it relies on a change in the physical state, cryo-neuromodulation poses no risk of changing the chemical or molecular composition of target neurons. Furthermore, although it cannot specify neuronal types, cannot be applied to neurons deep in the tissue, and has not been tested *in vivo*, it can potentially modulate any neurons on the brain surface, such as cortical neurons, without the need to wait for anesthetic chemicals to be absorbed^50^ or to be activated for specific ligand-gated ion channels^51, 52^. Finally, the small cooling effect area of our device allows highly localized modulation, which has been impossible with other cooling methods^8, 9, 10, 12, 53^. This allows single cell level study of cryo-modulation and the study of how it affects the network dynamics, offering faster and wider applications than chemical inhibitors, regardless of the neuron types.

Analyses of the amplitude and the width of spikes, ISI, and IBI show that cooling temperature slows down the action potential (AP) conduction velocities^29, 54, 55^, affects the regeneration of AP (Fig. 2d)^54^, decreases AP amplitude (Fig. 2e)^29^, and broadens AP width (Fig. 2l, m)^28, 56^. This may be a consequence of the direct inhibition of voltage-gated Na^+^ channels because the subcellular distribution of voltage-gated Na^+^ channels can change due to cooling^57^, interfering with AP generation. Furthermore, the resting membrane potential is depolarized at ∼ 15 °C, which may inactivate voltage-gated Na^+^ channels, slow the kinetics of voltage-activated Na+ currents^27^, reduce their amplitude, silence neuronal signaling, and ultimately delay recovery from cooling^43^ ^58, 59^ ^25, 33^. On the other hand, voltage-gated A-type K+ currents (IA currents), which are involved in regulating action potential shape, threshold, and inter-spike interval^30^, may also have been affected by cooling. Finally, K2P channels (TREK-1 and TRAAK) are highly temperature sensitive and can be significantly inhibited by cooling temperatures^60, 61, 62, 63, 64^. Overall, the effect of cooling may come from the physiological change of the channel dynamics at near 15 °C. Combined with our results about varying cooling duration in this study, we suggest that mammalian neurons can be silenced effectively by holding them at 15°C for one to five minutes.

Peltier^11, 12, 13, 14, 15, 65^, chamber^9, 10^, and helmet^53^ cooling devices have been used to study the effects of low temperatures on brain tissue, including the suppression of epileptic discharges^12^, the frequency changes of hippocampal theta rhythms^14^, and the song timing of bird brain in high vocal centre (HVC)^11, 13^, etc. Peltier-based probes have been used to cool specific brain regions as well^13, 14, 15, 16^. However, the cooling devices are either too large for localized cooling at the single-cell scale, or the probes are mostly thick. Long and Fee^13^ have used thickness of probe 330um, but they are slow to cool, unstable, and thawing is slower than cooling. A key discovery of our study is that the cooling is critical to recovering their activities after cooling. We found that neurons that were subject to slow cooling and stayed below 37°C for a longer period did not fully recover normal neural activities. On the other hand, when neurons were cooled fast with our device (at least 20℃/s), they fully recovered their normal activities.

In Fig 5, cooling of a limited area showed that the effects of cooling can be transmitted through the neuronal network. Cultured neurons show the limited axonal growth. When we observed the growth of axons under the microscope, it was difficult for axons to grow beyond 500 μm. However, there was cooling-induced silencing even at distances greater than 600 μm from the cooling point. Previous studies have shown that synaptic transmission is also sensitive to low temperature, which is consistent with our experimental findings^66, 67, 68^.

In this study, we developed a novel rapid cooling platform that reversibly inhibits neural activity. This cryo-neuromodulation system, which is based on the combination of a fast and spatially precise cooling device and MEA, is expected to serve as a new platform to study the effect of cryo-neuromodulation in neurons. Moreover, this platform can be combined with other chemical or optical neuromodulation techniques for more complex manipulation of neural activities, facilitating scientific discoveries. Finally, it offers a potential route to a new medical device for the precision-silencing of targeted neurons in e.g., epilepsy.

## Material & methods

### Configuration of cryo-neuromodulation system

The basical experimental equipment configuration of Cryo-neuromodulation system consists of a manipulator (PS7000C, Scientifica) which was combined with the cooling device, in vitro mcroelectrode array (MEA) systems (Multichannel systems, MEA2100) with the temperature controller enabling to maintain a temperature of 37 degrees, a head stage which can measure microcurrent and interface instrument connecting the head-stage with the temperature controller. Additionally, there is a microscope that was combined with CCD camera. The MEA (60MEA200/30iR-Ti-gr, Multichannel systems) has 59 electrodes fabricated by substance of titanium and gold. And the distance between each electrode is 200µm and the electrode of MEA can read the neural signal within 100 µm. The following, we decrease the temperature of the cooling device’s probe to using the Thermo electric cooling (TEC) controller (TEC-1123-HV-NTC-1M, Meerstetter engineering) and chiller (Thermo Scientific, Accel 500 LT). And it is used to supply 20℃ cooling water to cooling device to decrease temperature of cooling device to -10℃. (Suppelmentary Fig. 1)

### Rat primary cortical neuron culture

Primary cultures of rat cortical neurons were prepared from the brains of embryonic pups at day 18. First, pregnant female rats were sacrificed with CO_2_ and then abdominal skin was washed with 70% ethanol. After ethanol washing, abdominal skin was incised with surgical scissors. Through incised abdominal skin, day 18 embryos were taken out and transferred to 100Ø dishes with Hanks’ balanced salt solution HBSS (14185-052, Thermo Fisher Scientific) on ice. Before isolating brain, placenta was removed and then the heads were immediately cut through brain line by fine scissors. After getting rid of remaining embryonic skin and skull, the whole brain can be finely isolated. Set up cutting line in this step decides remaining non-brain part to be removed. From whole brain tissue, the cortex was dissected with fine surgical scissors. The meninges on dissected cortex was removed and then transferred to 15 ml tube. These cortex tissues were washed with fresh HBSS two times and then add 0.5ml of 0.05% Trypsin-EDTA (T/E) (25300-062, Thermo Fisher) and 0.5ml HBSS which make final concentration 0.025% into 15ml tube. Incubate the cortex tissues at 37℃, 5% CO2 for 10 min. After incubation, carefully remove the supernatant and avoid taking the remaining cortex tissues. Further washing step with HBSS two times and with medium two times. Finally, remove the supernatant and add fresh medium 1 ml. The cortex tissues were breakdown into single cells through pipetting. Following, we coated Microelectrode array (MEA) by Poly-L-Lysine (P2636, Sigma) and seeded neurons 5,000∼6,000 cells/mm2 on the NEA, put 1mL culture media. The Culture media consisted of Neurobasal medium (21103-049, Gibco), 2% B27 supplement (17504-044, Gibco), 2mM Gluta-max (35050-061, Gibco), 1% penicillin-streptomycin (15140-122, Gibco). Finally, In the condition of CO2 5% and 37℃, we incubated the MEA at least 14days. (Supplementary Fig. 2)

### Experimental procedures

The experimental procedures are as follows. First, we attached the MEA cultured neurons to the head stage of in vitro MEA systems and put CO_2_ chamber which can supply 5 % of CO2 on MEA and head-stage to minimize neurons damage before cooling by the cooling device. Then we measured normal signal of neurons. Next the CO_2_ chamber was removed before the temperature of the cooling device was lowered, since the cooling device interfered with the field of vision observing activated electrodes. To confirm the position of the probe of the cooling device and the activated electrode by the CCD camera which was combined the microscope, we moved the probe of the cooling device in x, y direction using the manipulator and finally adjust the z-axis position that was 50 to 100µm above bottom of MEA. Then, we decrease the temperature of the cooling device’s probe to using the TEC controller. And the experiment was conducted for cooling duration which were 1, 3, 5 and 10minutes at -10℃ and cooling velocity which were -20℃ per seconds and -4℃ per seconds. After cooling by the probe of the cooling device, we recovered the temperature of the cooling device’s probe to 37℃. Lastly, we analyzed the neural signals before, during, and after cooling, to understand the effect of cooling on neurons at each state.

### Data analysis of experiment

We selected the first spike in the burst from the measured neural signal by using the Neuro explorer (Version 5, Nex Technologies) program. Spike amplitude, spike shape, spike width values were obtained with sorted spikes. The average of the spike shape of each experiment was expressed in thin lines, and the mean of the thin line was expressed in bold lines. The standard error value of each experiment was expressed in a transparent surface. We categorized spike by using the online spike sorting program tridesclous (Samuel Garcia, Christophe Pouzat). As a result of spike sorting, we got 1 to 2 spike clusters. Also, there was no difference in the results of sorting the entire section and 3 section by divided into before, during and after cooling. Through the sorted spike, we obtained the values of inter-burst interval, inter-spike interval, spike rate, the number of spikes in burst, occurrence time of both first spike and recovery spike. Correlation matrix is based on spike rate histogram of spike timestamp.

### Analysis optimal condition to silence neurons

To observe neural signal silencing, we decreased temperature of cooling device. And The temperature is 37℃/s, 15℃/s, 0℃/s, -10℃/s. As a result, when the temperature of the cooling device reaches -10℃, neural signal was silence. The base for the result was as follows: When spike sorting was performed on the region that have been cooled, neural signal was not shown classified as spike but was perceived as noise. (fig. 1e). And we analyzed change ratio of spike rate before cooling and cooling. The calculation method of change ratio of spike rate is as follows. Change ratio of spike rate: [(Spike rate (after cooling) – Spike rate (before cooling)) / spike rate (after cooling)] * 100

### Analysis of neuronal signaling affected by cooling duration

We investigated how cooling duration affected neural signal. Cooling duration was 1, 3, 5minutes and 10minutes. And we compared the values of Inter spike interval (ISI), inter burst interval (IBI), spike width, spike amplitude, spike rate, occurrence time of first spike and recovery spike to analyze the effect of cooling on neural signal. Firstly, we classified the time of spike occurrence as first spike and recovery spike. Firstly, after the temperature was restored to 37℃, We called spike that was occurred for the first time ‘the first spike’. Following, we called as the recovery spike when the average number of spikes in a burst before cooling equals the average number of spikes in a burst after cooling. Following, the IBI times were not uniform in size of number, so these were rounded to represent natural numbers. And the results were shown in a histogram. In addition, the value of each cycle in the scatter plot in Fig. 2h was the average time of the IBI in each experiment. Next, the ISI average was calculated the average within the range of 0.01 ms to 0.8 ms. Finally, Spike width by cooling duration, normal line of gray color is average spike width of each experiments and bold line is spike width average of gray color lines. Also, transparent phase is standard error of spike width. Apart from that, the large neural signal generated at -10℃ was not the neural signal but the noise caused by impact on the culture media when the probe of cooling device was frozen or defrosted. (fig. 2a)

### Analysis of neural signal affected by cooling effect, depending on distance from cooling point

Neural signal in MEA randomly is generated. We wanted to do a quantitative analysis of neural signal affected by the cooling effect. So, areas on MEA were divided into four areas in a circular shape every 200 micrometers from cooling point. The quantitative analysis is changing ration of spike rate and silencing time of neural signal. And we defined that silencing time of neural signal is time of the last spike occurred after decreasing temperature to -10℃.

### Calcium imaging and observation

We prepared a stock solution by diluting dimethyl sulfoxide (DMSO) (D2650, Sigma) 1∼5 mM in the Fluo-4(F14201, Invitrogen) form of Powder, and kept it frozen at -20℃. When experiment was conducted, working solution was made mix 2 µL of Fluo-4 into 2 ml of HBSS. The preparation process for calcium imaging was as follows. We removed the culture media from the cultured neuron on 13Ø petri dish and clean it with HBSS 3 times. Then the working solution is put in a 13Ø petri dish as much as amount of the culture media you put in when the neuron cultured. Next, we put the petri dish in the incubator and incubated it for 30minutes. After finishing incubation, we replaced the solution of the Incubated petri dish to HBSS 2∼3 times. The calcium images are monitored using an inverted microscope (IX71, Olympus) with a CCD camera (True Chrome 2, TUCSEN). To analyze the intensity of calcium image quantitatively, photobleaching-guideline was defined as the brightness of neurons steadily was decrease without changing intensity by photo-bleaching phenomenon. Analysis of neural signal affected by cooling velocity of cooling device The experiment was conducted at -20℃/s and -4℃/s to investigate the effect of the cooling velocity on the neural signal. The time to reach -10℃ for each velocity was about 2.4 seconds and 12 seconds (See supplemental). We maintained temperature for 2 minutes at 37℃/s. And it cooled 1 minute based on when the temperature began to decrease. We repeated that. we called 37℃ ‘before cooling’, and after cooling, 37℃ regions were named ‘trial #’. Particularly, the local field potential was shown as significant change as soon as the cooling started from the cooling device with cooling velocity -20℃/s of cooling device. It could be assumed for neurons to have been greatly stimulated by the cooling. (fig. 2a-b) On the other hand, when cooling velocity of the cooling device was -4℃/s, the local field potential didn’t be shown as significant change.

### 2-sample Kolmogorov-Smirnov test

The KS test is one of the methods of comparing the two samples because they are sensitive to differences in location and shape of the empirical cumulative distribution function. In analysis of this experiment, we used the kstest2 function of Matlab (R2019b, Mathworks). ‘Data from x1 and x2 are derived from the same continuous distribution for vector x1, x2’ returns the test results for null hypothesis and the alternative is ‘x1 and x2 are derived from different continuous distributions’. If the test rejects the null hypothesis at the 5% significance level, the result *h* is 1 but opposite case is 0. Based on this, we tested whether the distribution before and after cooling is the same for cooling duration and cooling velocity experiments through two-sample test.

### Local cooling device construction and operation

To perform the in vivo electrophysiology experiments in neuronal networks in this work, the cooling setup requires not only controllability in both target temperature and the rates of cooling and thawing but also the capability of localized cooling. For these requirements, we developed probe cooling device, for which a tungsten tip with 200 µm diameter connected to a copper plate cooled by two Peltier units to locally cool a certain part of the neuronal network. The cooling system can produce a target temperature down to -20 °C and its maximum cooling rate is up to -20°C /s. The tungsten cold probe was electrically insulated from the electric circuits that powered Peltier modules, so that delicate electric signals from the neuronal network could be simultaneously measured by MEA system during the probe cooling process without cross-noise. Fig. 2c-d shows the illustrative description of our cooling system: a tungsten probe that provides cooling to the neurons, a copper plate that cools the tungsten probe, two Peltier modules that cools the copper plate, and two heatsinks that cools the two Peltier modules. Given that the tungsten probe required to submerge in the solution during cooling, its side wall was thermally insulated with a polymer jacket. The two Peltier modules were controlled by a PID controller. The tungsten probe was set to have a distance 90µm above from the neurons and localized cooling was provided.

### Simulation analysis for cryo-neuromodulation experiment

Because the temperature at the location of neurons could not be directly measured with any currently available temperature sensor, we calculated the temperature at the neurons based on known thermal properties of solution, tungsten, and polymer jacket. COMSOL Multiphysics® was used to calculate the neuron temperature, which incorporated Navier-Stock’s equation, kinetic equation, and thermal energy equation. The grid independence test was performed that no dependence of the calculated results on the mesh size was observed. The boundary conditions were set to reflect the actual experimental conditions that the temperature at the bottom of MEA substrate was maintained at 37°C and ambient temperature was set to be 20 °C with 5 W/m^2^-K convection coefficient. Three isothermal surface conditions at the copper plate were used, 15 °C, 0 °C, -10 °C, and -20 °C. The boundary surface between liquid and solid is applied to non-slip condition, and the liquid is supposed to incompressible laminar flow. As shown in Fig. 2e, the temperature at the neurons is calculated to be 12.6 °C for the copper plate temperature of -20°C, which linearly increased with the copper plate temperature.

## Data availability

The data are available from the corresponding author upon reasonable request.

## Acknowledgments

This work is supported by a National Research Foundation of Korea (NRF) grant (2020R1A4A2002728, 2020R1A2C2009093, 2021R1A2C3004262) funded by the Korean government (MSIP), and also supported by Kinship Foundation, Alfred Sloan Foundation, and Klingenstein-Simons Foundation.

## Author contributions

JK and JP designed the experiment. JK performed the experiments and analyzed the data. JSL isolated neurons. SN contributed to installing cooling device and designing simulation. ES contributed to carrying out the experiments. JL contributed to configurating experiments. TK contributed to design experiment. SWC, GK, SSK, and JP supervised and led the research. JK, SSK and JP wrote the manuscript. All authors discussed the results and contributed to the manuscript.

## Additional information

**Supplementary information** is available for this paper at https://doi.org/

**Correspondence** and requests for materials should be addressed to Seung-Woo Cho, Gunho Kim, Sung Soo Kim, or Jungyul Park.

**Reprints and permission information** is available at http://www.nature.com/reprints

**Publisher’s note** Springer Nature remains neutral with regard to jurisdictional claims in published maps and institutional affiliations.

## Supplementary Information

**Supplementary Fig. 1.**
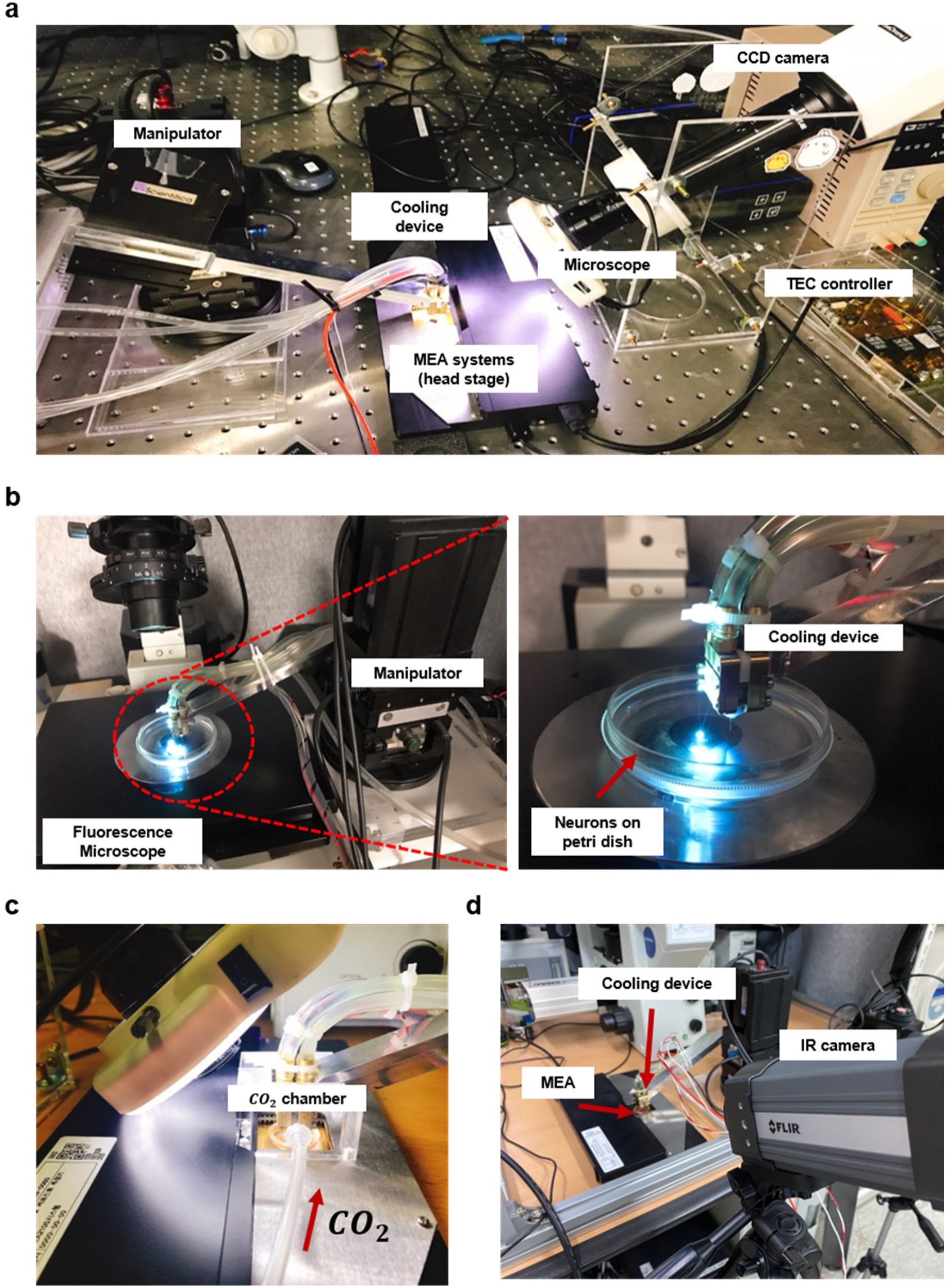
Configuration of cryo-neuromodulation equipment for each experiment. (a) Configuration of equipment for cooling duration experiment, cooling effect by distance from cooling point, and cooling velocity experiment. (b) Left: For calcium imaging experiment, configuration of equipment. Right: Magnified cooling region. (c) Mounted CO_2_ Chamber on Head stage of MEA systems (d) Configuration of media’s surface temperature measurement experiment with IR camera.

**Supplementary Fig. 2.**
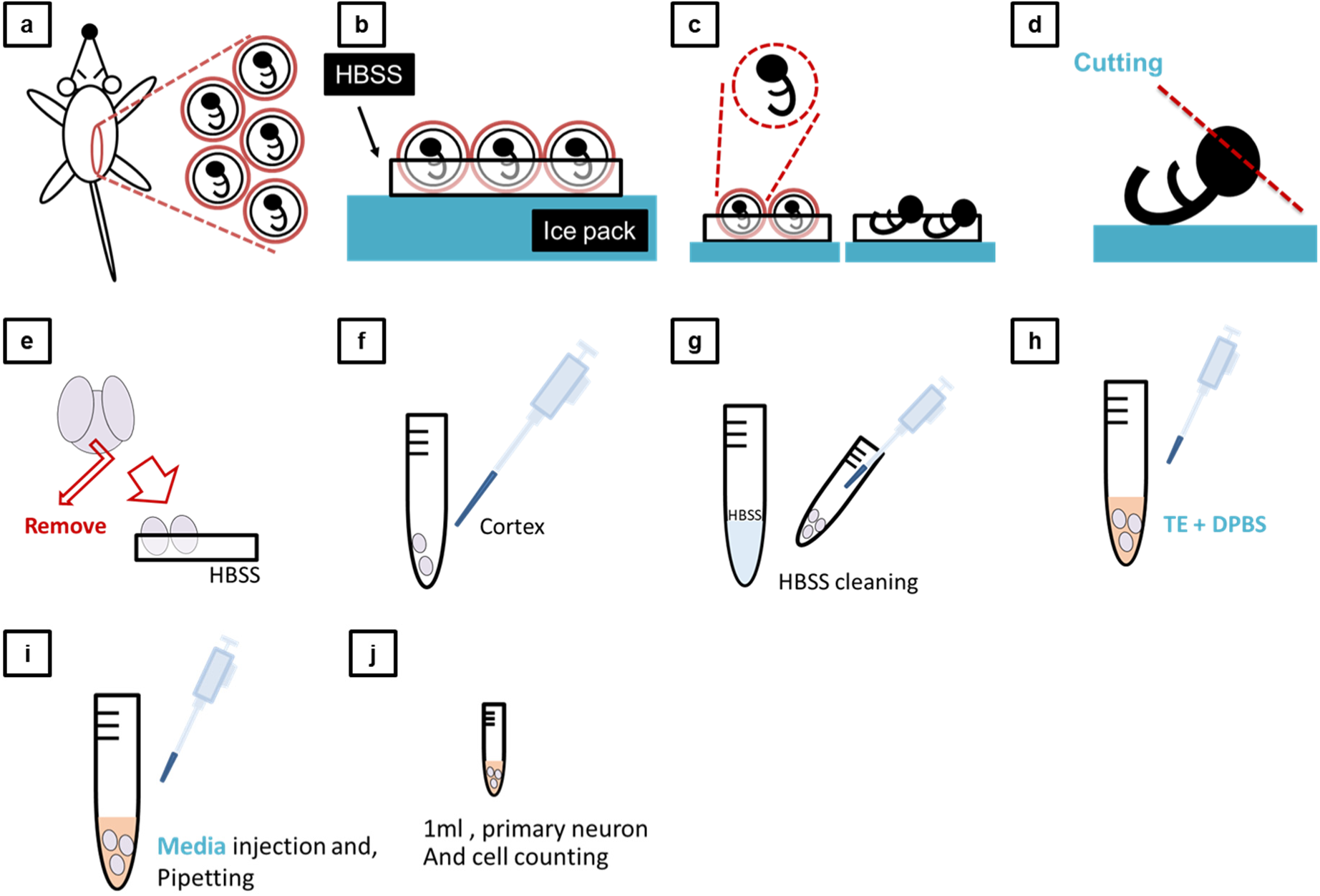
Isolation neuron. (a) Pregnant female mice at gestational day 18 were subjected to abdominal dissection for fetal extraction. (b) The extracted fetuses were immediately immersed in Hanks’ Balanced Salt Solution (HBSS) or Dulbecco’s Phosphate-Buffered Saline (DPBS) to maintain physiological conditions. (c) The placenta was removed from the fetuses to facilitate subsequent brain isolation. (d) The fetal head was excised, and the brain was carefully extracted for further processing. (e) The cortex was isolated from the extracted brain and immersed in HBSS or DPBS for additional purification. (f, g) The isolated cortex was transferred to a 15 ml tube, where cleaning procedures were performed. (h) Subsequently, 0.05% Trypsin-EDTA (TE) and DPBS were added to the tube, and the cortex was incubated for 5-10 minutes in an incubator to facilitate enzymatic digestion. (i) Following the incubation period, culture media were injected into the tube, and the cortex was mechanically dissociated into a single-cell suspension through pipetting. (j) The final cell suspension was subjected to cell counting procedures, ensuring accurate quantification.

**Supplementary Fig. 3.**
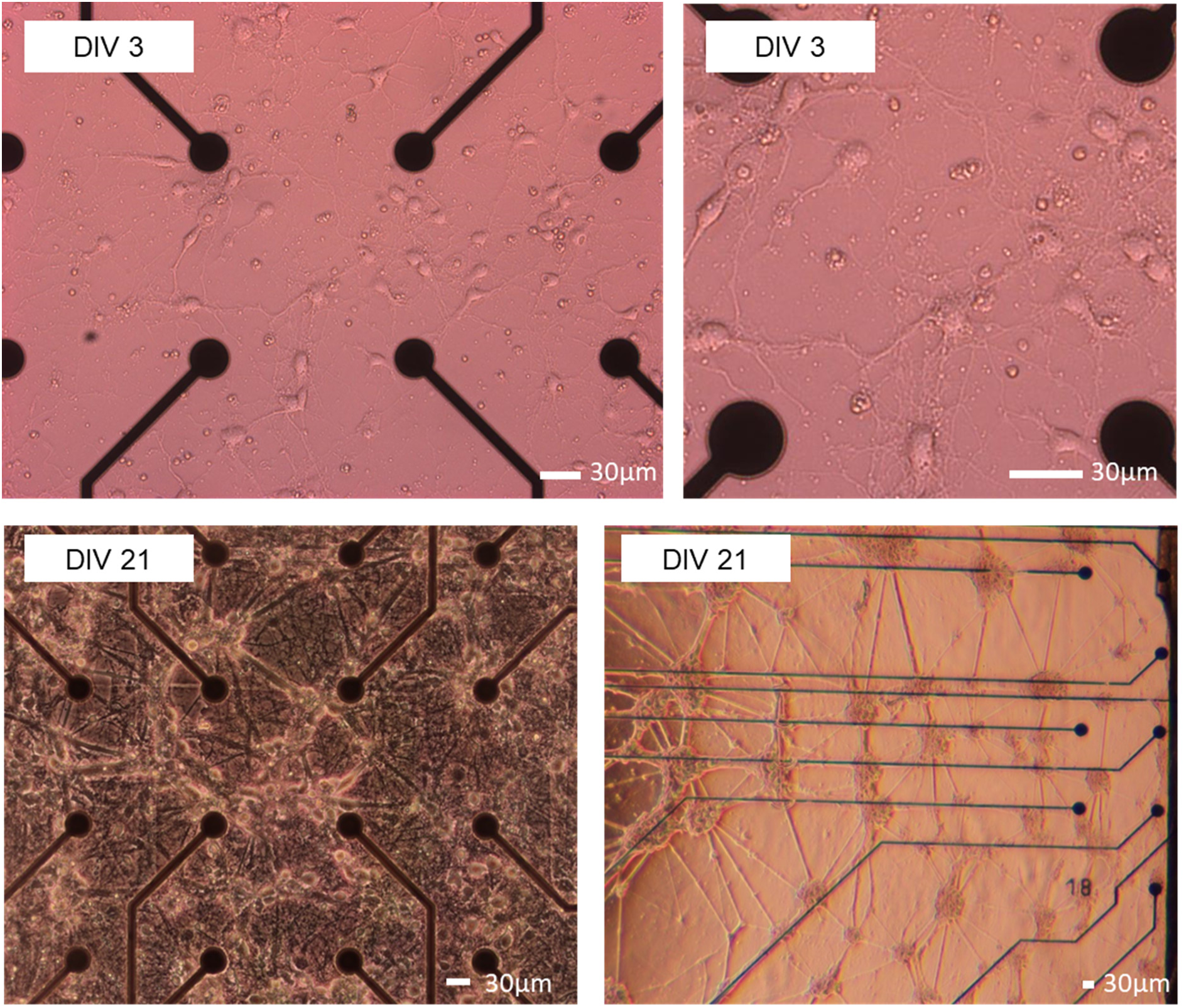
Microscope image of neuron. Top: Day in vitro (DIV) 3 neurons, Down: DIV 21 neurons

**Supplementary Fig. 4.**
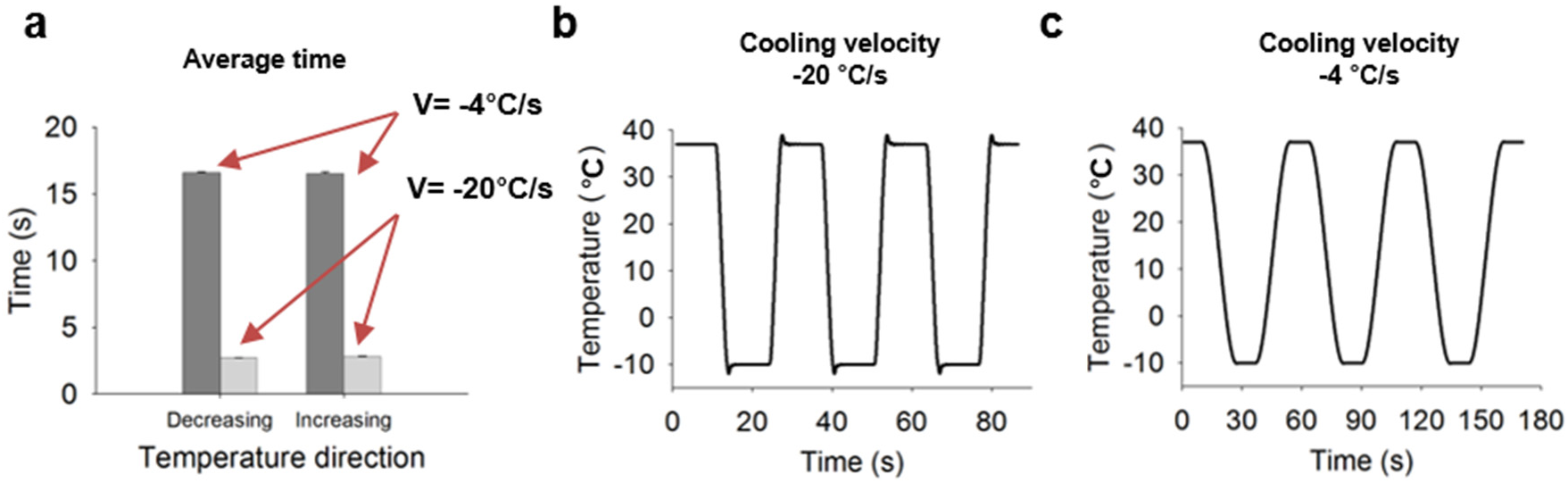
Performance of cooling device by cooling velocity. (a) Average time of reaching target temperature by each cooling velocity. (b) When cooling velocity is -20°C/s, temperature graph of cooling device from 10°C to 37°C. (c) When cooling velocity is -4°C/s, temperature graph of cooling device from 10°C to 37°C.

**Supplementary Fig. 5.**
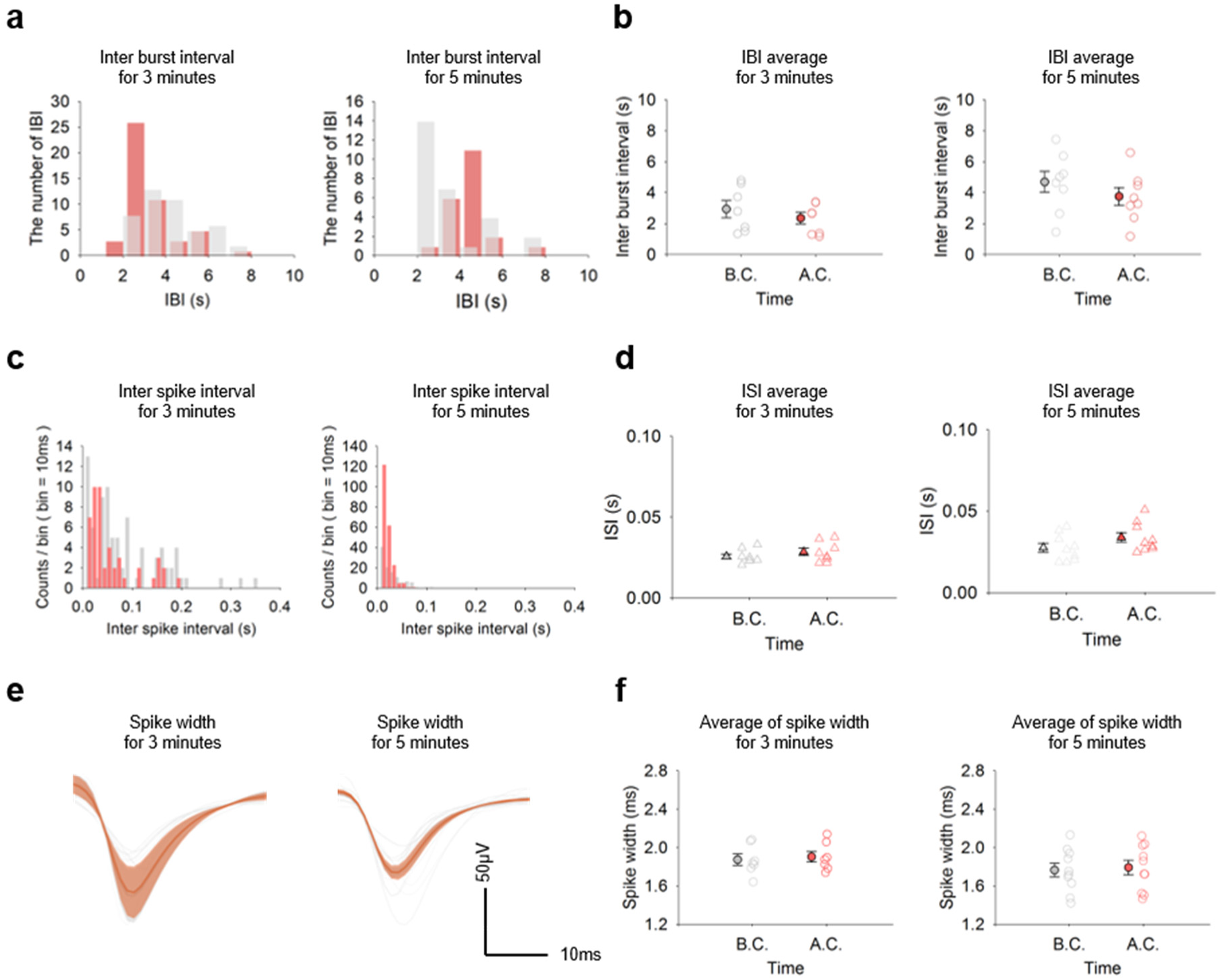
The recovery of neural activity depends on the duration of cooling at 3, 5 minutes. (a) Left: The number of IBI histogram at 3-minutes Right: the number of IBI histogram at 5-minutes. (b) Scatter plot of inter-burst interval before and after cooling. Left: At 3-minutes (paired t-test: Before cooling vs After cooling, n=8, p=0.5189). Right: At 5-minutes minutes (paired t-test: Before cooling vs After cooling, n=9, p=0.6030). (c) Inter-spike interval histogram; Left: At 3-minutes. Right: At 5-minute. (d) Scatter plot of ISI’s average Left: At 3-minute (paired t-test: Before cooling vs After cooling, n=8, p=0.9290). Right: At 5-minutes (paired t-test: Before cooling vs After cooling, n=9, p=0.2500). (e) Average and Standard error spike width. (see Methods) Left: At 1-minute. Right: At 10-minutes. (f) Scatter plot for average of spike width before and after cooling Left: At 3-minute (paired t-test: Before cooling vs after cooling, n=8, p=0.9290) Right: At 5-minutes (paired t-test: Before cooling vs after cooling, n=9, p value=0.6030).

**Supplementary Fig. 6.**
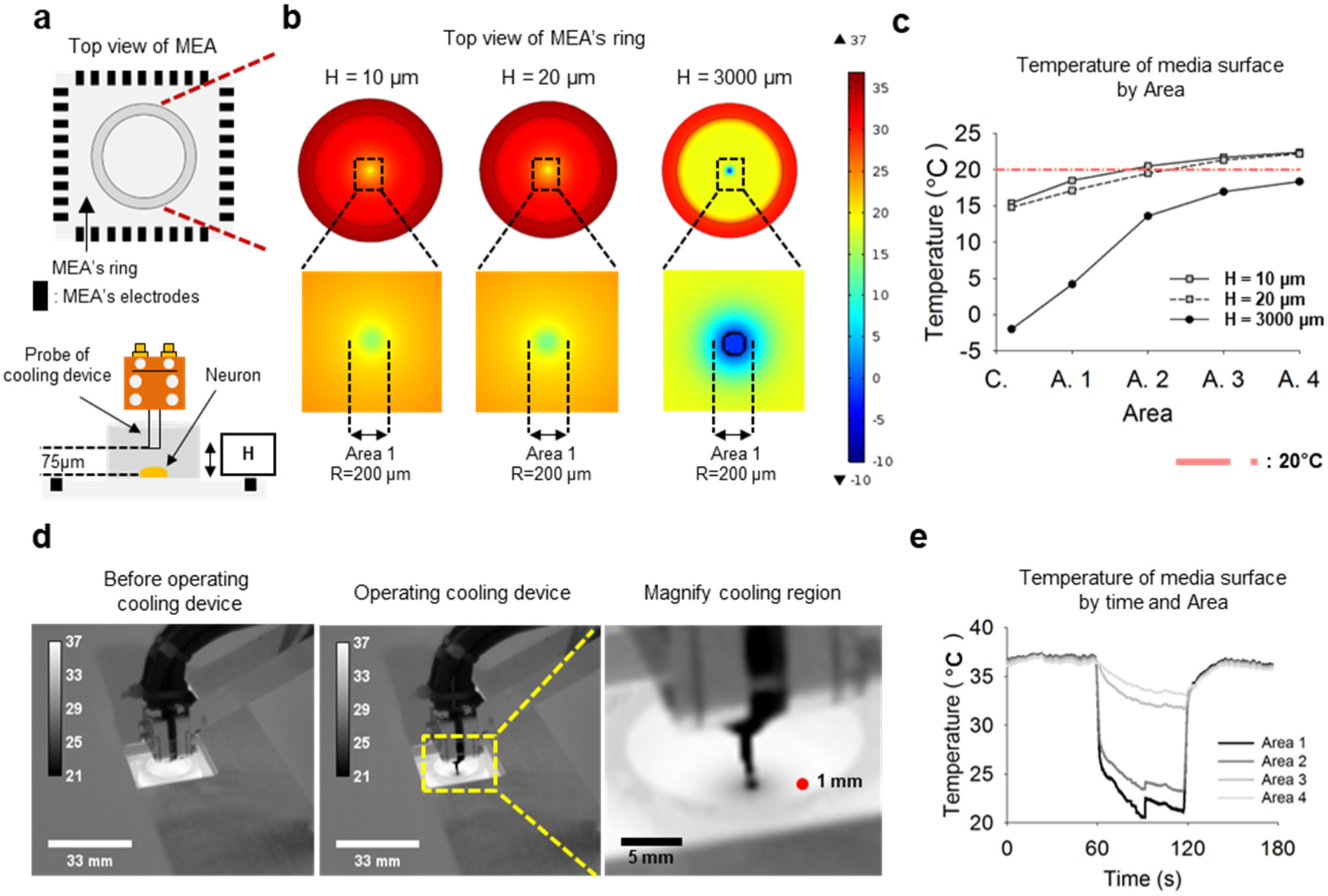
Work to know actual temperature for neurons. (a) Top: top view of MEA. Down: side view of MEA, H is height of culture media (b) Right: When H is 10 µm, temperature distribution on culture media by simulation results. Center: H is 20 µm. Left: H is 3000 µm. (c) Result in heat transfer simulation; temperature of media surface by Area#. (See Extended Data Fig. 4 in detail) (d) Left: Monitoring cooling region with IR camera at 37℃. Center: Left: Monitoring cooling region with IR camera at 10℃. Right: Magnify cooling region at 10℃. (e) Surface temperature graph of culture media at cooling duration 1 minute.

**Supplementary Fig. 7-1.**
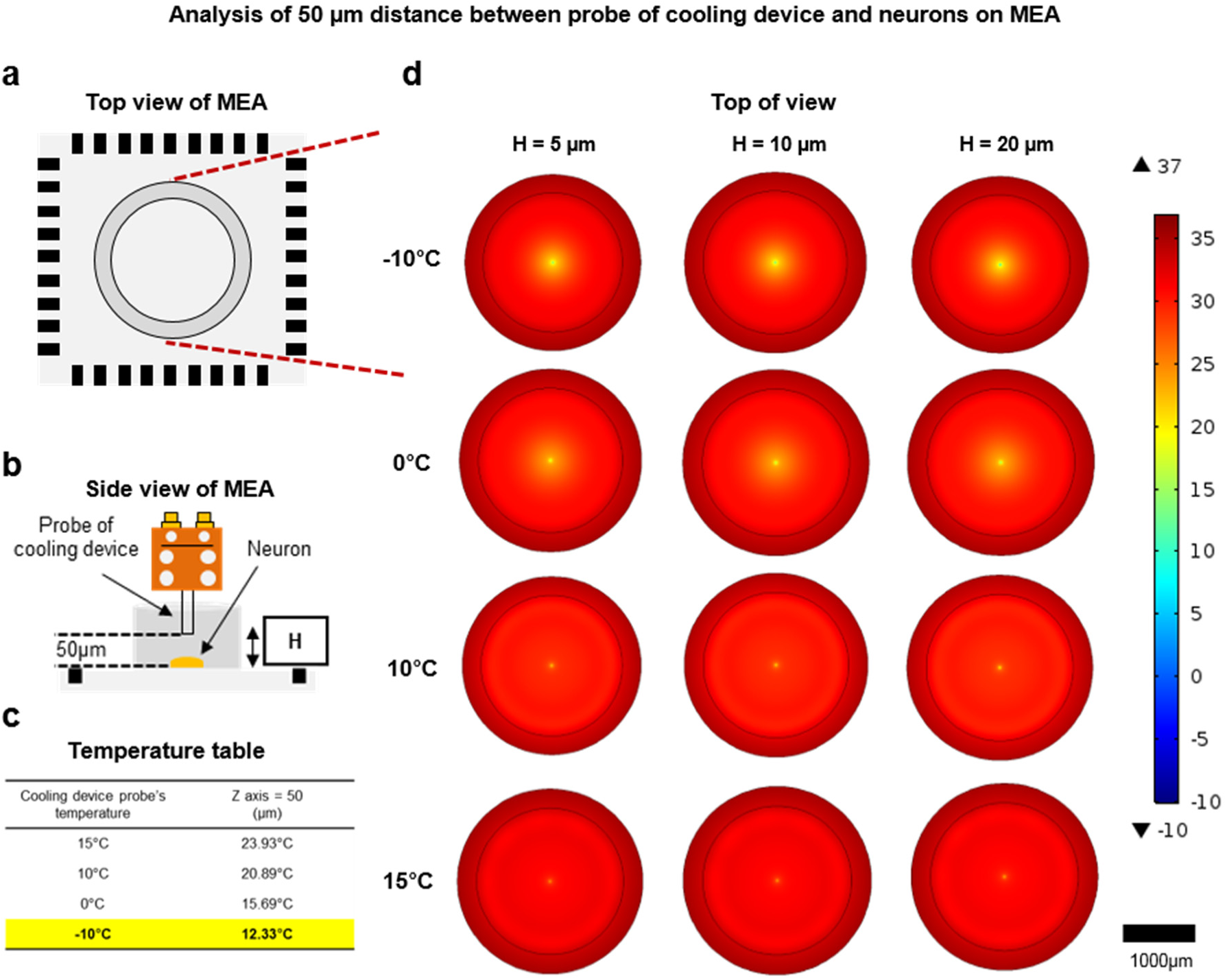
Heat transfer simulation to know actual temperature for neurons. Analysis of 50µm distance between probe of cooling device and neurons on MEA. (a) Top view of MEA. (b) Side view of MEA and probe of cooling device. (c) When 50µm distance between probe of cooling device and neurons on MEA, temperature of cooling device and actual temperature for neuron with simulation. (d) Result in heat transfer simulation by culture-media height and temperature of cooling device.

**Supplementary Fig. 7-2.**
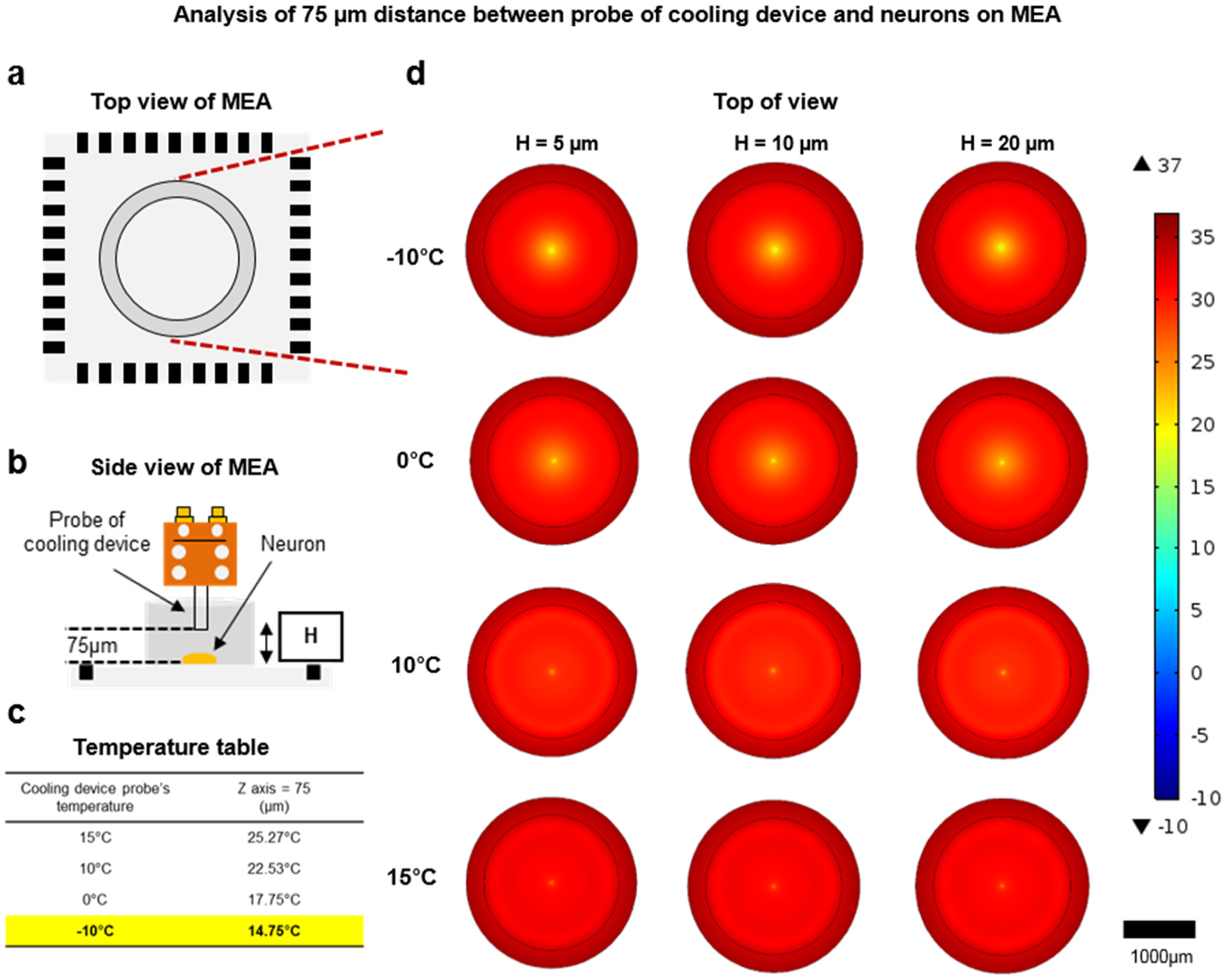
Heat transfer simulation to know actual temperature for neurons. Analysis of 75µm distance between probe of cooling device and neurons on MEA. (a) Top view of MEA. (b) Side view of MEA. (c) When 75µm distance between probe of cooling device and neurons on MEA, temperature of cooling device and actual temperature for neuron with simulation. (d) Result in heat transfer simulation by culture-media height and temperature of cooling device.

**Supplementary Fig. 7-3.**
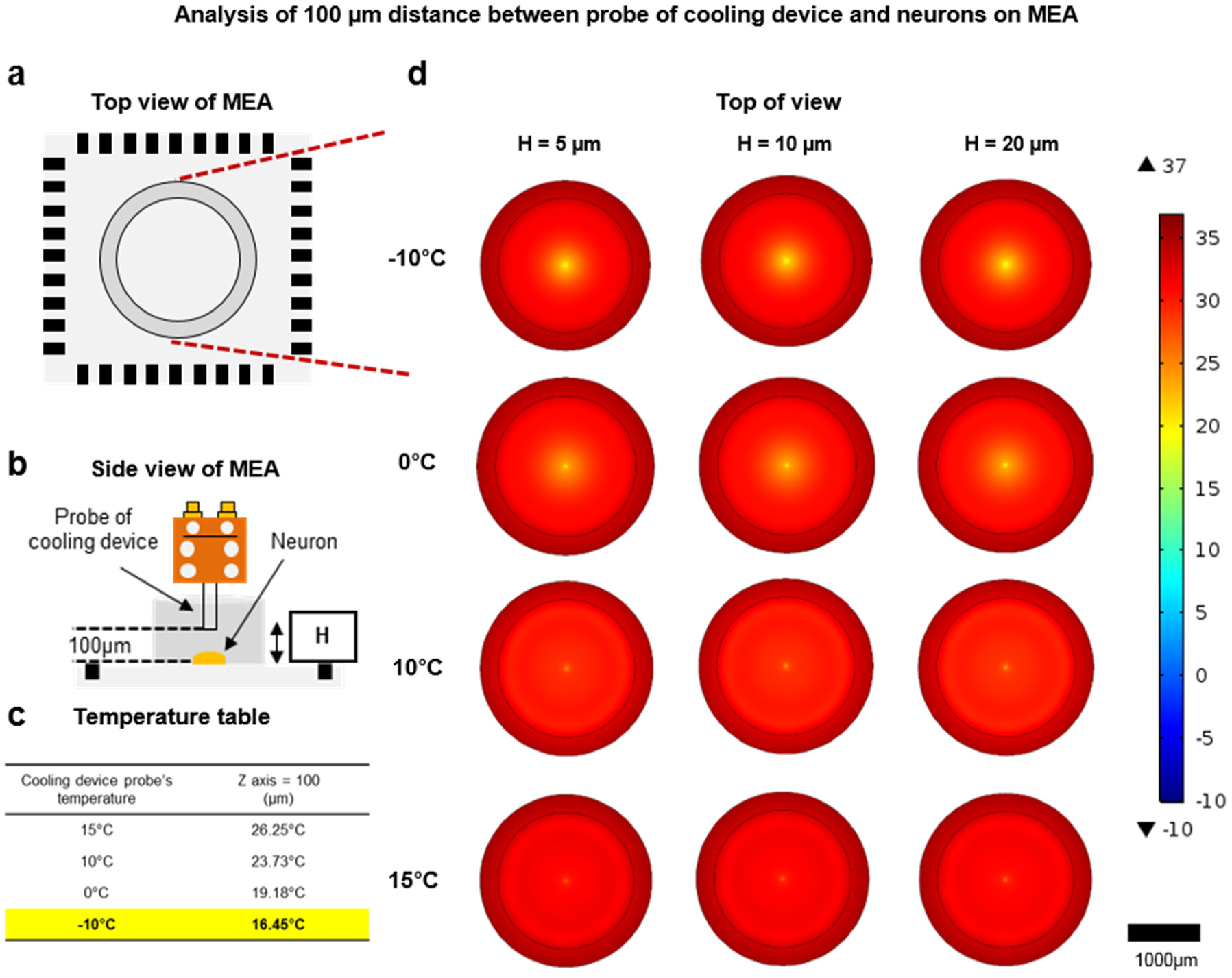
Heat transfer simulation to know actual temperature for neurons. Analysis of 100µm distance between probe of cooling device and neurons on MEA. (a) Top view of MEA. (b) Side view of MEA. (c) When 100µm distance between probe of cooling device and neurons on MEA, temperature of cooling device and actual temperature for neuron with simulation. (d) Result in heat transfer simulation by culture-media height and temperature of cooling device.

**Supplementary Fig. 8.**
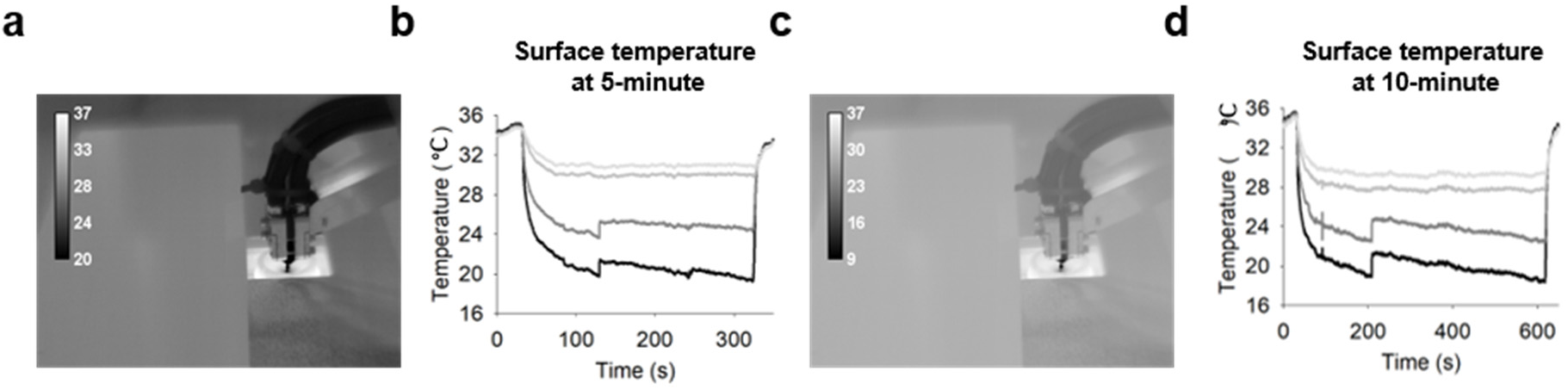
When cooling, temperature observation of culture-media surface at 5, 10 minutes. (a) Monitoring cooling region with IR camera at cooling duration 5-minutes. (b) Surface temperature graph of culture-media surface at cooling duration 5-minutes. (c) Monitoring cooling region with IR camera at cooling duration 10-minutes. (d) Surface temperature graph of culture-media at cooling duration 10-minutes.

**Supplementary Fig. 9.**
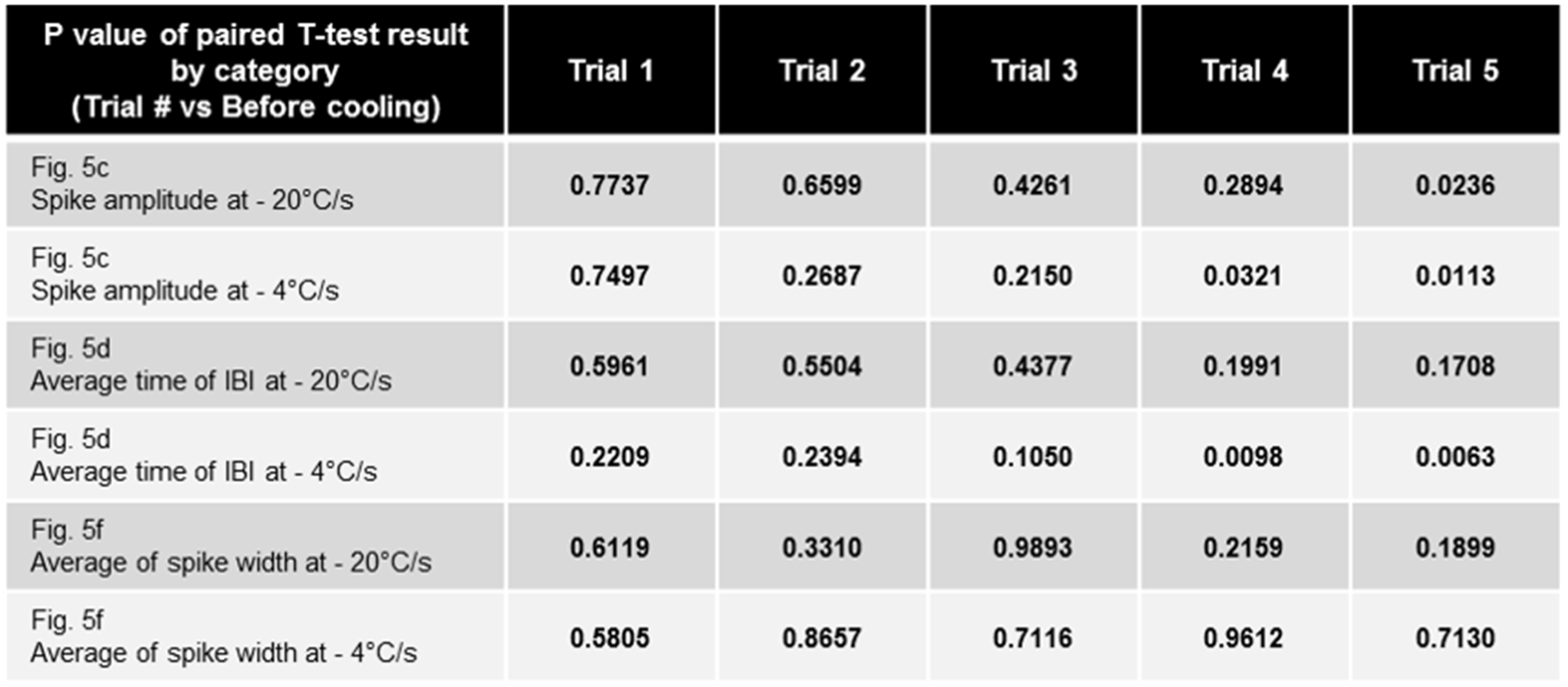
P values table of paired T-test result by Figure 5.

**Supplementary Fig. 10.**
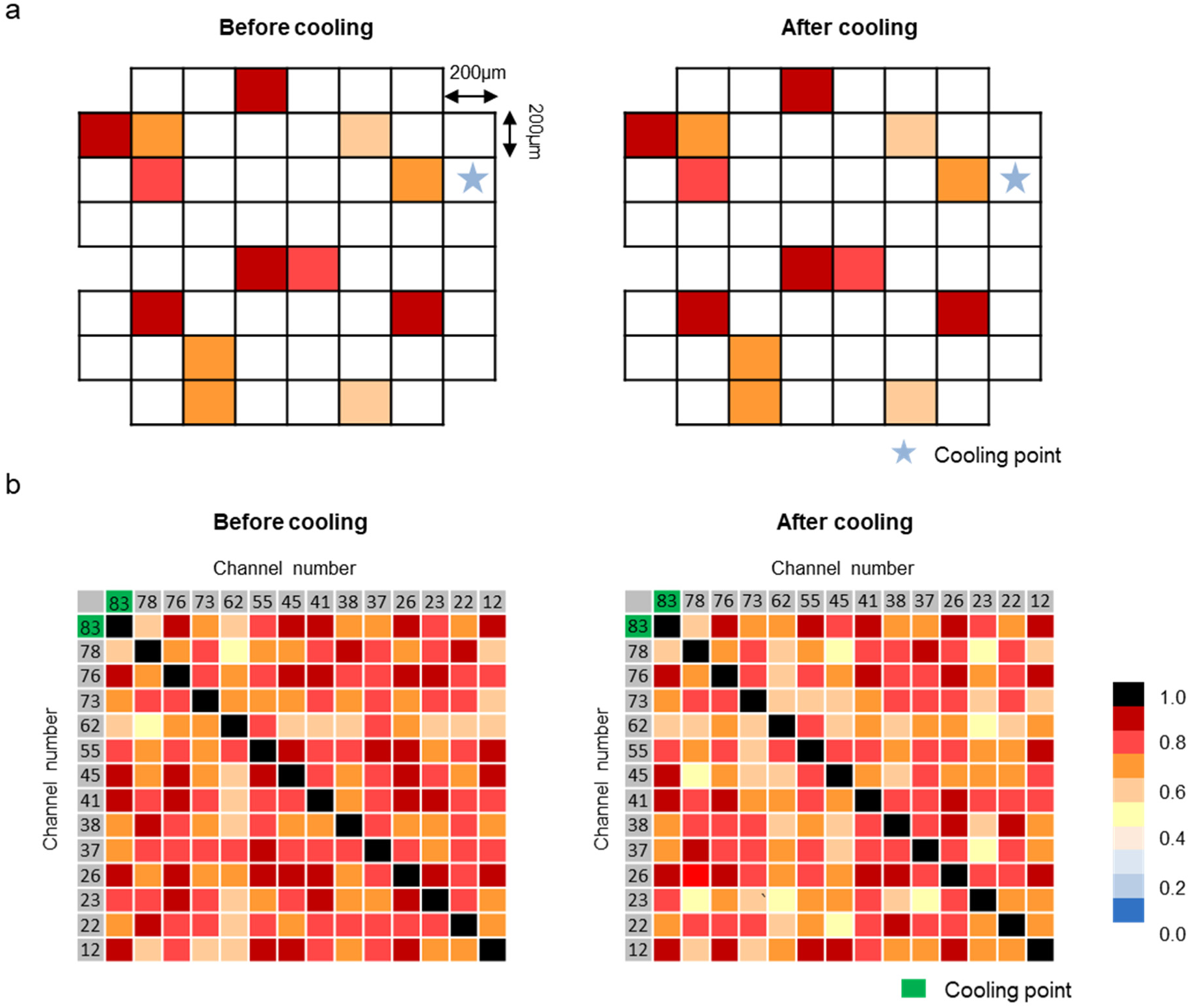
Correlation matrix in the case that all active channels were silenced. (a) Correlation matrix between cooling point and other active channels before and after cooling. White cells indicate inactive channel. (b) Correlation matrix between active channels before and after cooling.

**Supplementary Fig. 11.**
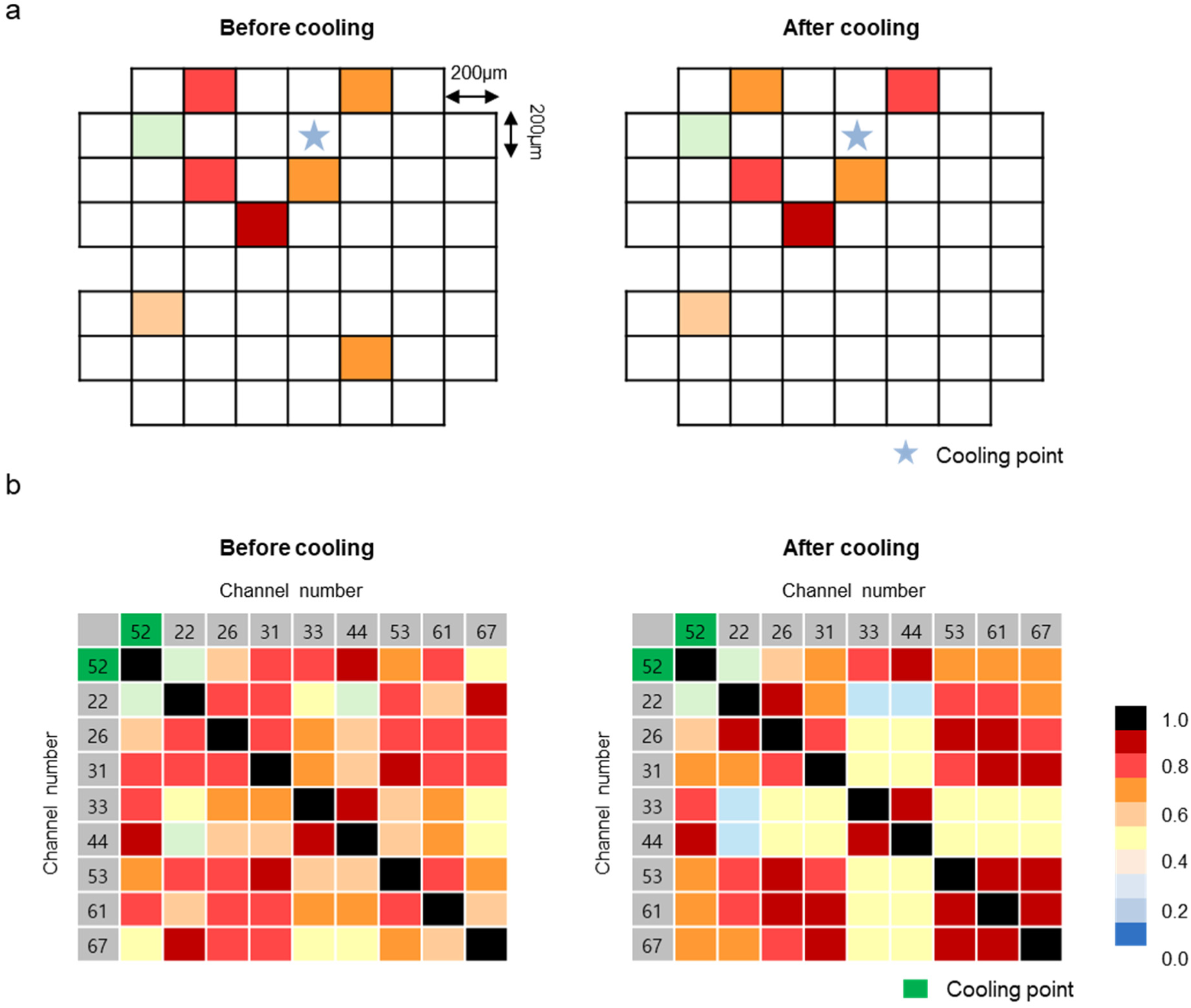
Correlation matrix in the case that a portion of active channels were silenced. (a) Correlation matrix between cooling point and other active channels before and after cooling. White cells indicate inactive channel. (b) Correlation matrix between active channels before and after cooling.

